# The genetic robustness of RNA and protein from evolutionary, structural and functional perspectives

**DOI:** 10.1101/480087

**Authors:** Dorien S. Coray, Nellie Sibaeva, Stephanie McGimpsey, Paul P. Gardner

## Abstract

The reactions of functional molecules like proteins and RNAs to mutation affect both host cell viability and biomolecular evolution. These molecules are considered robust if function is maintained despite mutations. Proteins and RNAs have different structural and functional characteristics that affect their robustness, and to date, comparisons between them have been theoretical. In this work, we test the relative mutational robustness of RNA and protein pairs using three approaches: evolutionary, structural, and functional. We compare the nucleotide diversities of functional RNAs with those of matched proteins. Across different levels of conservation, we found the nucleotide-level variations between the biomolecules largely overlapped, with proteins generally supporting more variation than matched RNAs. We then directly tested the robustness of the protein and RNA pairs with *in vitro* and *in silico* mutagenesis of their respective genes. The *in silico* experiments showed that proteins and RNAs reacted similarly to point mutations and insertions or deletions, yet proteins are slightly more robust on average than RNAs. *In vitro*, mutated fluorescent RNAs retained greater levels of function than the proteins. Overall this suggests that proteins and RNAs have remarkably similar degrees of robustness, with the average protein having moderately higher robustness than RNA as a group.

**Significance Statement:** The ability of proteins and non-coding RNAs to maintain function despite mutations in their respective genes is known as mutational robustness. Robustness impacts how molecules maintain and change phenotypes, which has a bearing on the evolution and the origin of life as well as influencing modern biotechnology. Both protein and RNA have mechanisms that allow them to absorb DNA-level changes. Proteins have a redundant genetic code and non-coding RNAs can maintain structure and function through flexible base-pairing possibilities. The few theoretical treatments comparing protein and RNA robustness differ in their conclusions. In this experimental comparison of protein and RNA, we find that they have remarkably similar degrees of overall genetic robustness.

## Introduction

RNA is mainly known for its role in translation, yet it is also involved in controlling gene expression (e.g., bacterial small RNAs and microRNAs), intrinsic immunity (e.g., CRISPR-mediated acquired immunity) and the cell’s response to environmental stimuli (e.g., thermosensors and riboswitches) (1, 2). The discovery of functional non-coding RNAs (ncRNAs), like transfer RNA and catalytic RNAs, led to the proposal of an ancestral RNA world, where RNA both catalyzes the reactions of life and encodes genetic information (3). Despite the importance of non-coding RNAs to cellular function, many RNAs exhibit low sequence conservation and are not as broadly distributed as their proteinaceous brethren (4). In contrast, proteins make up the bulk of the biomolecular contents of the cell, and are required for the majority of critical cellular structures and functions. Yet protein production is a complex, multi-stage process that is sensitive to errors (5, 6).

The ability to preserve a phenotype in the face of sequence perturbations is termed mutational robustness (7, 8). More robust molecules maintain their phenotypes despite mutations, while less robust molecules lose their function rapidly. The genotype level at which mutations are considered, and the types of phenotypes (structure, function) that are measured, can vary between analyses. Here, we consider mutations in nucleotide sequence and how they modify phenotypes such as the structure and function of encoded genes. The robustness of individual protein and RNA molecules are both affected by factors such as the stability of individual molecules, interactions and gene expression levels (9–13). For example, RNA stems are generally more robust than loops (9, 10) and allow non-canonical base-pairs (14) while protein alpha helices are more robust than beta strands and both were more robust than unstructured coils (11, 12). Proteins and RNAs also have independent mechanisms that allow for near-neutral changes that maintain molecular structures and functions (Table 1).

**Table 1:**
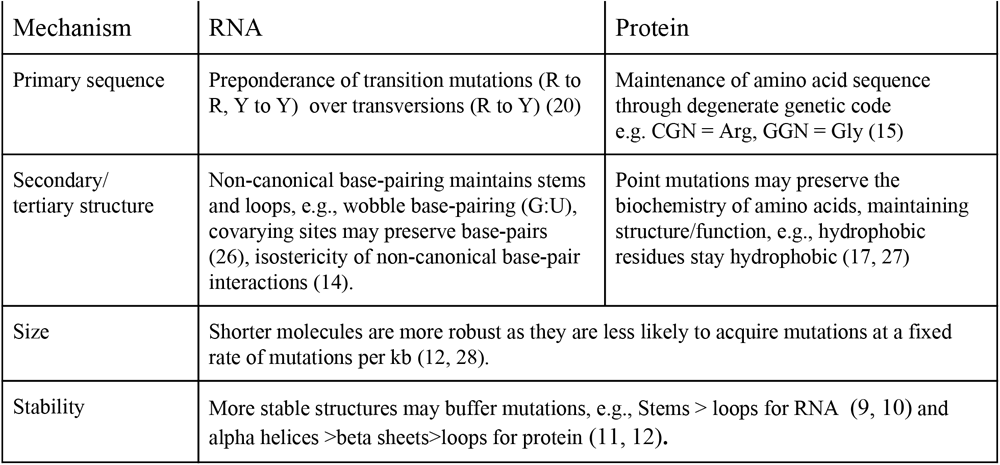
Avenues of neutral change within RNAs and proteins (20, 25).

Protein robustness relies, in part, on the robustness of the genetic code. Degeneracy of the genetic code allows for mapping of up to six codons to the same amino acid with interchangeable tRNAs (15, 16). Furthermore, when point mutations change the amino acid, the new amino acid coded for is likely to have similar biochemical properties due to the code’s organization (17). Monte Carlo simulations have shown that the extant genetic code is significantly more robust to substitution and frameshift mutations than randomly generated genetic codes (18–22). Premature stop codons and frameshift errors can be introduced by substitutions or indels (insertions and deletions). While ncRNA production requires transcription—and in some cases, additional maturation such as editing, splicing or processing—proteins require these in addition to translation, which depends upon the maintenance of a correct reading frame (23). These additional steps likely amplify the potential harm of nucleotide changes (5). This is supported by the fact that disease-associated sequence variation is enriched ten-fold in human protein-coding regions (24) and that overall variation is reduced in coding regions, particularly indels (6). Because of this, we hypothesize that RNAs are more robust to mutation than proteins, and can tolerate greater sequence change while maintaining function.

Previous comparisons of protein and RNA have involved computational analysis of neutral networks: a collection of related sequences that give rise to the same phenotype. Earlier analysis using reduced genetic codes (e.g., G+C for RNA and hydrophobic:hydrophilic for protein) (20, 29–32), found that RNA networks differed from protein networks, being larger (more robust) but also less compact. More recent work has shown that this is dependent on the mathematical framework used, and suggested that proteins and RNAs are more similar (31, 32).

To compare protein and RNA robustness on as fairly as possible we have designed a series of experiments that explore the question of which is more robust. Given the variety of mechanisms and levels at which molecules can alter robustness (summarized in Table 1) and the biochemical differences between RNAs and proteins, it is impossible to make any truly fair comparisons. Therefore we have carefully selected matched pairs of proteins and RNAs that share a function or structure, and compare their evolutionary conservation over matched time periods, their structural robustness with simulated mutations and their functional robustness with error-prone PCR. In exploring the broad differences between protein and RNA robustness under comparable conditions, we may gain insight into the origins and bioengineering potential of protein and RNA.

## Results

We have devised three tests, two *in silico* and one *in vitro*, to explore whether protein and RNA differ in their robustness to mutation. **1.** We have considered the degree of DNA sequence variation between pairs of matched RNA and protein families over matched time-periods. To control variation as much as possible, ncRNA genes were compared to the genes of proteins that are involved in related functions (e.g., ribosomal RNAs and ribosomal proteins, riboswitches and the proteins these regulate, tRNAs and tRNA synthetases, etc.). These protein-RNA pairs have shared recent phylogenetic and selection histories. Our expectation is that more robust genes will tolerate more mutations over fixed timescales, and thus exhibit greater sequence change than less robust genes (see Figure 1). **2.** We have simulated point mutations and insertions/deletions in DNA sequences encoding interacting proteins and ncRNAs (ribonucleoparticles/RNPs) with solved tertiary structures. The potential impacts of mutations have then been assessed based upon structural stability (ΔΔ G), evolutionary profiles (Δ bitscore) and predicted secondary structure profiles (see Figure 2). **3.** We have compared the functional robustness of a protein and RNA directly with mutagenesis of a functionally comparable fluorescent RNA (33, 34) and fluorescent protein (35–37), and quantified the fraction of mutants that maintained function for each class of molecule (see Figure 3).

**Figure 1:**
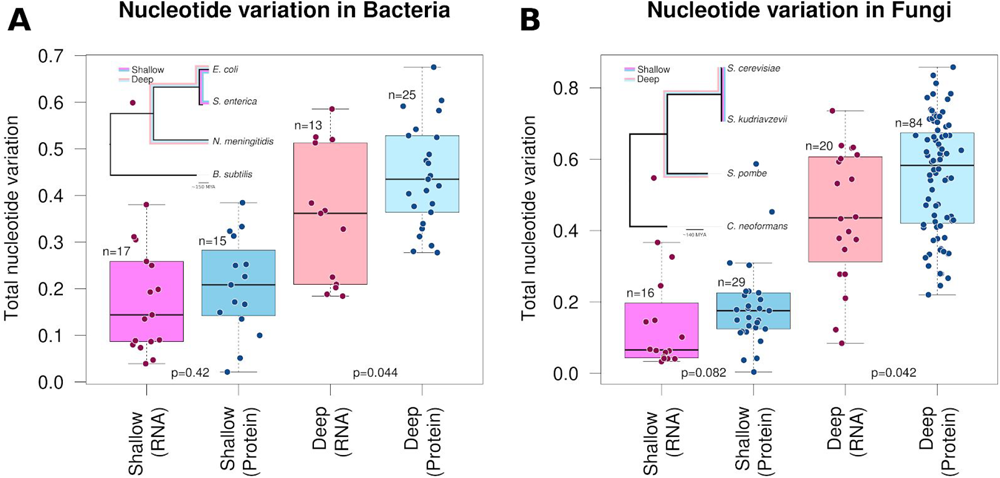
Proportion of variable nucleotides between shallow and deep divergence times for conserved protein and RNA families. The proportion of total nucleotide variation was calculated for aligned orthologous proteins (blue) and RNAs (pink) and illustrated with box-whisker and jitter-plots. (**A**) Proteins and RNAs from representative bacteria, *Escherichia coli* and *Neisseria meningitidis* (deep divergence, lighter shades) or *E. coli* and *Salmonella enterica* (shallow divergence, darker shades). **(B**) Orthologous protein and RNA families from representative fungi, *Saccharomyces cerevisiae* and *Schizosaccharymocyes pombe* (deep divergence, lighter shades) or *S. cerevisiae* and *Saccharomyces kudriavzevii* (shallow divergence, darker shades). **Tree inserts:** The relationship between the deep (lighter shades) and shallow (darker shades) diverged species on SSU rRNA, (dnaml) (38), phylogenetic trees. It should be noted that the ‘shallow’ species are estimated to have diverged less than 150 million years ago (MYA) in each case, whereas the ‘deep’ species diverged approximately >400 million MYA in each case.

**Figure 2:**
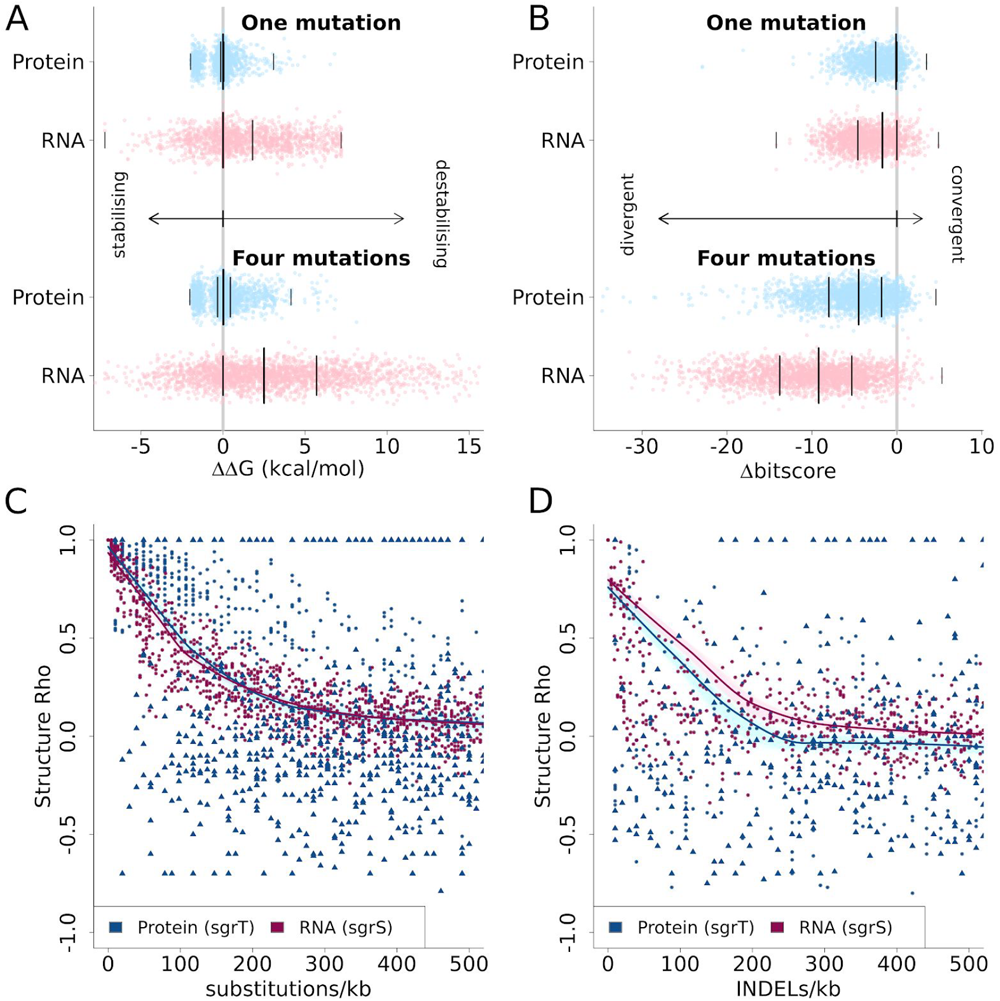
Robustness of structure predictions to random *in silico* mutagenesis for selected protein-RNA pairs. (**A&B**) For 24 non-redundant pairs of RNP structures available in the PDB, either one (upper) or four (lower) point mutations have been randomly introduced into corresponding DNA sequences. The impact of these simulated mutations on protein and RNA sequences have been assessed using predicted Δ bitscore (**A**) and ΔΔ G (**B**) values. The distributions of values are illustrated using jitter-plots, with the medians, 25^th^ and 75^th^ percentiles, and extremes indicated with a vertical line for each distribution. (**C&D**) Comparing the robustness of protein and RNA secondary structures to random substitutions or insertion/deletion (INDEL) mutations. Secondary structure probabilities were predicted for native and mutated sequences, and the per-residue probabilities of alpha/beta/coil structures (protein) or base-paired/not-base-paired (RNA) were compared between native and mutated sequences using Spearman’s correlation (Structure Rho). Truncated proteins or sRNAs with a length less than 75% of the original are indicated with a solid triangle, otherwise a solid circle is used. Truncated regions and other unalignable residues were excluded from the correlation calculation. The average trends between mutation rates and Structure Rho are indicated with local polynomial regression (loess) curves. To indicate the confidence for each loess curve, these were bootstrapped 500 times and plotted in light pink (RNA) or blue (protein) to resampled points.

**Figure 3:**
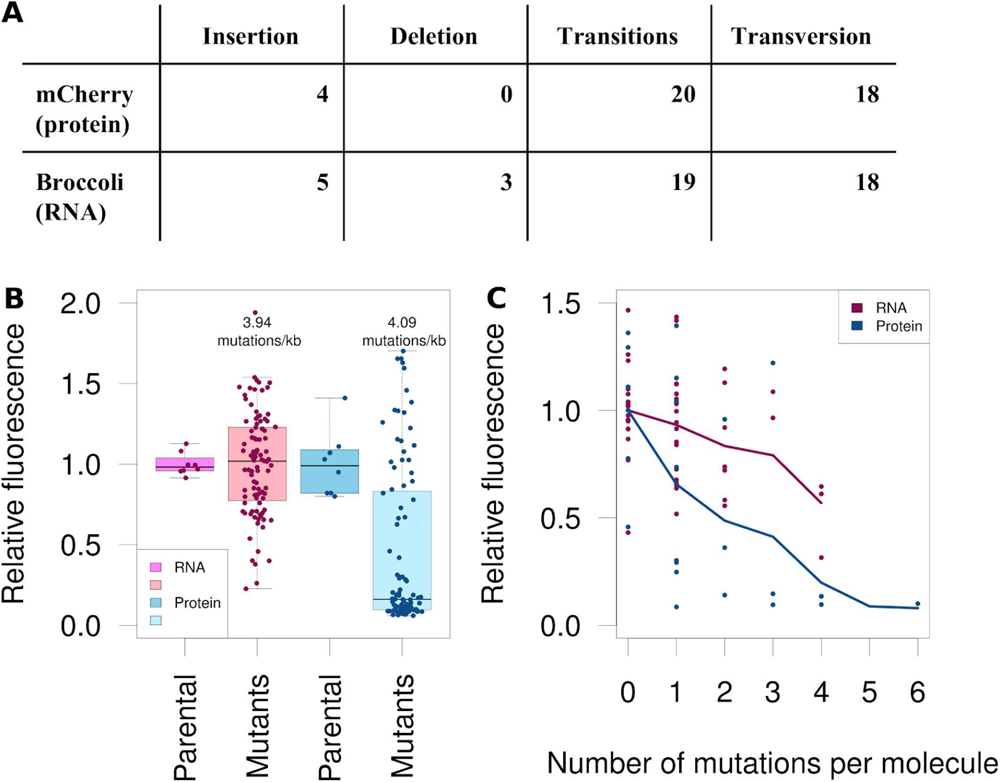
Relative fluorescence intensities of mutated RNA Broccoli and mutated protein mCherry. Libraries of randomly mutated fluorescent RNA aptamer Broccoli and fluorescent protein mCherry have been generated and tested for function relative to an unmutated control. (A) The mCherry and Broccoli libraries were matched for similar rates of mutations per kilobase (kb) (4.09 and 3.94, respectively) using an error-prone PCR protocol. The fluorescence intensities for 96 mutants each of the RNA and protein were compared with those for eight unmutated controls. Measurements were recorded for three separate replicates. (B) Individual molecules of mCherry and Broccoli mutants were sequenced and their fluorescence compared using the number of mutations per molecule (zero to six). (C) We counted the different types of variants that were observed in the sequenced mutants.

### Experiment 1: Sequence diversities of ncRNAs and proteins involved in the same processes

Our expectation is that nucleotide variation in proteins and RNAs between diverged species will indicate the degree of neutral variation that has occurred while the gene functions have been preserved. More robust genes are expected to tolerate more mutations over time. We have collected Rfam RNA and Pfam protein-domain pairs that are involved in the same biological processes, and are found in the Figure 1 reference genomes (39, 40). These can be broadly classified as ribonucleotide particles (RNPs), cis-regulatory elements and corresponding proteins, and dual-function genes where one partner is modified or processed by the other (Tables S1 & S2). These RNA and protein pairs have similar selection histories because of their shared functions, though their individual structural and/or catalytic constraints will vary.

We curated **four** independent sets of RNA:protein pairs in yeast and bacteria species that were divergent enough to exhibit sequence diversity and have matched G+C contents (~50%) (Figure 1). Two sets are for “deep” gene pairs that are highly conserved, often involved in functions such as transcription or translation. The species compared have diverged more than 400 million years ago (*Saccharomyces cerevisiae* and *Schizosaccharymocyes pombe* for yeast, *Escherichia coli* and *Neisseria meningitidis* for bacteria). The other two sets are for “shallow” gene pairs that have diverged recently. These gene-pairs generally have a limited phylogenetic distributions, and possibly a high tolerance of mutations that is a characteristic of new genes (41, 42), which are often involved in functions such as gene regulation, biosynthesis and maturation of RNPs. The species compared have diverged less than 150 million years ago (*S. cerevisiae* and *Saccharomyces kudriavzevii* for yeast, *E. coli* and *Salmonella enterica* for bacteria).

At each level of conservation, we computed the total number of variable nucleotide sites between the two species for a given RNA and protein domain. This was normalized by the length of the sequence to derive the total nucleotide variation for each. Variants were further classified as neutral or not, depending on whether secondary structure (RNA), amino acids (protein) and/or the biochemistry of the RNA or protein was preserved (Figure S1).

Generally, the level of total nucleotide variation within protein-coding sequences was higher than the RNAs, yet the distributions for both largely overlapped. Only the distributions for just the deep gene pairs could be considered to be statistically significantly different (p=0.044 in bacteria, and p=0.042 in fungi, two-sided Wilcoxon rank-sum test) (Figure 1A&B). While the shallow gene pairs were not statistically significantly different from each other (p=0.42 in bacteria, and p=0.082 in fungi, two-sided Wilcoxon rank-sum test) (Figure 1A&B).

It is possible that interactions between the RNAs and proteins constrained the degree of variation between the two, with one of the pair evolving slower because it maintained interactions with a slowly evolving partner (43). We did not, however, see a general significant correlation between the rate of nucleotide variation in a given RNA and its matched protein (Figure S2), leading us to conclude that this was most likely not the case.

The sequence diversity between populations provides an indication that a gene’s function is robust to changes in the nucleotide sequence. While protein-coding mRNAs tend to have greater nucleotide diversity than RNAs it is typically not significantly higher over short time-scales, yet is marginally significant over longer timescales. By this measure, RNAs and proteins exhibit comparable degrees of robustness.

### Experiment 2: simulated mutations and the predicted impacts on structural and evolutionary robustness

We have simulated point mutations and insertions/deletions in a number of selected pairs of protein and RNA encoding DNA sequences. The impacts of simulated mutations have been analysed using tools for assessing mutation impacts on structure or function that work on both protein and RNA.

For the first simulation we use non-redundant RNPs with solved structures (e.g. tRNA-synthetase pairs, signal recognition particles, 6S RNA bound by RNA polymerase, components of the spliceosome, a guide RNA in a complex with a Cas protein, and snoRNAs with associated proteins) available in the PDB (July 2019) (44). We have inferred the theoretical impact of point mutations on structures by estimating changes in Gibbs free energy, (ΔΔ G, units: kcals/mol) (45, 46) (Figure 2B). The variants have been further analysed using an orthogonal approach for assessing evolutionary robustness using differences in profile HMM scores (Δ bitscore, units: bits) (47, 48) (Figure 2A). Each mutant sequence could be classed as divergent from the parental (zero) if the corresponding Δ bitscore value is less than zero, or as destabilised if the corresponding ΔΔ G is greater than zero. We found that the proportion of ncRNA mutants that were classed as either divergent or destabilised was consistently greater than that for proteins (see Table S1 for the numerical values).

In addition to the above, we have conducted a more detailed analysis of secondary structure robustness on a dual-function protein/ncRNA pair SgrS/SgrT, where the correlation between predicted probabilities of secondary structure elements have been compared between native and mutated sequences (49). SgrS and SgrT were selected as they are both structured and short (for computational efficiency) and the protein and RNA structures are not contained in the Protein Data Bank (PDB) that the secondary structure prediction tools we use were trained on (e.g. PSSpred).

We compute per-residue secondary structure probabilities on native simulated point or insertion/deletion mutant sequences. The correlation between native and mutant per-residue secondary structure probabilities is computed (“Structure Rho” i.e. Spearman’s correlation coefficient) and the distributions of these for a range of mutation densities have been plotted (Figure 2C&D). Both the protein and RNA mutants retain about half the parental structure (structure correlation of 0.5) at 100 point mutations per kilobase, though the protein shows a higher variance in response to mutation compared to RNA (Figure 2C&D). The protein is slightly more sensitive to indels than RNA, but shows a very similar overall level of decline in its structure.

### Experiment 3: Mutational robustness of a functionally equivalent RNA and protein

It is possible that the structure could be maintained but function lost, or that some molecules may continue to function better than others despite changes in structure (i.e., they are more robust). To test for robustness of function, we have mutated an RNA and a protein matched for an assayable function (fluorescence) and tested these mutations *in vivo*.

To investigate how biomolecules may differ in their robustness to mutations in DNA, we constructed mutant libraries of the fluorescent RNA aptamer Broccoli (33, 34) and the fluorescent protein mCherry (37). Both these molecules have been developed synthetically in the laboratory, and have not been subjected to strong evolutionary pressure outside of fluorescence. With a mutation frequency of approximately four mutations per kilobase, the relative fluorescence intensity for the population of Broccoli mutants was significantly (*P* = 1.5 × 10^−13^, Wilcoxon rank-sum test) more than that for mCherry (Figure 1A). Though the median fluorescence of the Broccoli population decreased slightly as the frequency of mutations increased, even at six mutations per kilobase, the Broccoli library had higher relative fluorescence intensities than the mCherry library with four mutations per kilobase (Figure S3). At 234 bases, the gene for Broccoli is much shorter than that for mCherry (711 bp). We sequenced approximately 40 molecules from each library and compared the number of mutations per molecule. Broccoli retained more of its fluorescence than mCherry with the same amount of mutations per molecule (Figure 1B). The frequencies of different types of mutations that occurred in the biomolecules are similar, with few insertions or deletions (indels) and similar numbers of transitions and transversions.

## Discussion

We have hypothesized that RNA would be more robust than proteins. This is supported by the fact that RNA often requires less processing than protein to produce functional molecules (50), is not susceptible to frameshift mutations (51) and is less likely to be found over broad evolutionary distances with homology searches (possibly due to high higher mutation rates) (4). Our multi-scale tests of RNA and protein robustness revealed no consistent evidence to support our hypothesis. This leads us to conclude that both molecules are remarkably robust to mutation, and that proteins are likely to be slightly more robust than RNAs based upon the balance of all the evidence. This finding corroborates some previous theoretical studies that suggest proteins and RNAs have similar overall robustness to mutation (31, 32). Our investigation of RNA and protein robustness was, in part, initiated by the observation that RNAs and proteins are differentially distributed across phylogenetic distances (4). A possible explanation for the difference is a higher mutation rate for functional RNAs, making them difficult to detect. However, we did not find any evidence to support this possibility when comparing the total nucleotide variation of matched RNAs and proteins over matched evolutionary divergences. If RNAs are not more robust than proteins, as our experiments imply, other factors must account for the apparent differences in phylogenetic distributions. For example, it is likely that protein homology search is statistically more powerful than that for nucleotides (52, 53). It could also be speculated that gene turnover and neofunctionalization are more rapid for RNAs than for proteins (42).

If RNAs were more robust than proteins, we would expect phylogenetically and functionally matched RNA families to have more nucleotide diversity than proteins of the same evolutionary background. Comparing the sequence diversity of extant RNAs and proteins revealed similar patterns in bacteria and fungi. Rates of nucleotide substitution were close, with proteins generally having more nucleotide diversity than RNAs. This was also seen when comparing Δ bitscore and ΔΔ G of RNPs mutated *in silico*. The proportion of divergent/destabilised ncRNAs was similar but higher than that seen with proteins. A specific protein-RNA pair, SgrS/SgrT was tested more in depth to both point mutations and indels. The RNA was slightly more robust than proteins in response to indels, however the overall responses were similar in both molecule types. These measures of RNPs are proxies of robustness, but taken together suggest that RNA is no more robust than proteins. They responded remarkably similarly but protein appears to be somewhat more robust than RNA.

The genetic robustness of proteins, especially to indels, is unexpected for a few reasons. The need for translation increases the avenues of error in protein production, and the evolution of translational robustness has been considered a factor in constraining nucleotide diversity in highly expressed proteins (13). While INDELs can greatly change downstream amino acids through frameshifts, RNA has no code to protect, and can, in theory, absorb additional nucleotides in stem bulges (9, 54, 55). Nonetheless, the predicted structures for RNA SgrS were almost as sensitive to indels as the protein SgrT.

The functional test of the fluorescent RNA Broccoli and protein mCherry was the only test where the RNA was considerably more robust. In order to match the mRNA and ncRNA sequence lengths as closely as possible we used a double Broccoli aptamer, which means a disabling mutation in one half can leave the other half with functionality. Consequently, we may have inadvertently increased the robustness of the RNA over the protein.

Future evolutionary, theoretical and experimental studies are needed to explore the initial results presented here. We acknowledge the limitations of using just one RNA and protein pair for analysis, the mutagenesis could be repeated with different fluorescent RNAs and phylogenetically distinct fluorescent proteins or other comparable functions such as self-splicing introns and inteins. Furthermore, simulated evolution experiments such as and directed evolution could be performed to identify differences between protein and RNA evolvability and robustness (56). Alternatively, experimental evolution datasets could be mined for appreciable differences between protein and RNA evolution (57). Robustness and evolvability may also be modeled computationally using flow reactor simulations (58). Forms of robustness other than mutational robustness, like the robustness of protein and RNA interaction networks, and robustness to environmental conditions such as temperature (59), pressure and pH fluctuations (60) can also be explored. The study of the interplay between robustness and evolvability informs our understanding of how new functions evolve in proteins and RNAs (31, 61–63), the proposed transition from an RNA world (64) and will help biomolecular engineering of functions under mutational and environmental challenges.

## Methods

### Natural variation of functional ncRNA:protein systems

We have curated pairs of RNA and protein families from Rfam and Pfam that are linked by either direct interactions or by process, and conserved between either *E. coli* and *N. meningitidis* (deep divergence; n=26 pairs) or *E. coli* and *S. enterica* (shallow divergence; n=18 pairs) in bacteria; in fungi we investigated RNPs from, *Saccharomyces cerevisiae* and *Schizosaccharymocyes pombe* (deep divergence; n=98 pairs) or *S. cerevisiae* and *Saccharomyces kudriavzevii* (shallow divergence; n=49 pairs). The RNA and protein coding sequences were aligned using the most accurate techniques appropriate for each molecule type (53, 65). Each pair of deep or shallow diverged nucleotide sequences were aligned, for RNA using Rfam, tRNAscan-SE (v1.3.1) or Intron covariance models (66–68) and `cmalign` (v1.1.1) (69) or, for the protein domains, using `hmmalign` (v3.1b2) (70) and concordant codon-aware nucleotide alignments generated with `PAL2NAL` (v14) (71). The total number of variant sites was recorded for each alignment, and a nonparametric Mann-Whitney U rank-based test (72) implemented in R was used to compare the distributions of the number of variant sites for protein and RNA. The results can be viewed in Figure 1 and Supplementary Tables 1&2.

The phylogenetic trees embedded in Figures 1A&B are crafted from SSU rRNA alignments. Sequence alignments for the three model bacterial or fungal species and an outgroup were generated by aligning to the corresponding Rfam covariance model (73) (bacterial SSU - RF00177 and eukaryotic SSU - RF01960, for the fungi) with `cmalign` (69), a phylogenetic tree was estimated for each alignment using `dnaml` (v3.69) from the phylip package (38, 74) with default parameters.

The corresponding computer code for the above steps is available in ‘`computeSynonNonsynon.pl`’ and ‘`plotConservation.R`’ scripts in the github repository. Raw data is available in the `data/genomes-bacterial/` and the `data/genomes-fungal/` directories.

### Simulated variation and ribonucleoprotein secondary structure

A curated list of RNP structures was selected by analysing the RNA sequences available in the PDB sequence repository ftp://ftp.wwpdb.org/pub/pdb/derived_data/pdb_seqres.txt.gz -- (18 July 2019). RNA sequences were extracted and checked for plausible minimum free energy structures using `RNAfold` from the Vienna RNA Package (75). RNP selections were based upon the presence of a stable RNA structure and diversity (e.g. avoiding an over-abundance of any particular structure type, or species). In addition, ncRNA lengths between 50 and 150 nucleotides, and protein lengths between 50 and 500 were favoured (although some exceptions have been made).

Messenger RNAs for the selected proteins were recovered by `tblastn` search of the NR database. If necessary, mRNAs were manually corrected to match the corresponding PDB sequences. Quality control steps included verifying that RNA, protein and mRNA sequences were consistent with the corresponding PDB entries. Protein and RNA sequences were further checked for corresponding profile models in the EggNOG database (v5.0) (76) and Rfam (v14.0) (73) using HMMER3 profile HMMs (70) and Infernal covariance models (69) respectively. See Table S3 for a list of the final 24 structures that met our criteria.

In order to compute ΔΔ G values (change in Δ G - Gibbs free energy - due to mutations), we selected a machine-learning method, `MAESTRO` (77) for the proteins, and a nearest-neighbour free-energy method, `RNAfold` (75, 78) for the RNAs. Missing values due to method failures were recorded as NA’s in our output files. `MAESTRO` failed on 15% of the four mutations, and 4% of the one mutation simulations, while `RNAfold` failed on 0.2% of the four mutation simulations. In the worst case, an additional 15% high ΔΔ G values for proteins would not have altered our conclusions. In order to compute Δ bitscore values (change in profile HMM or CM bitscores due to mutations (47)), we used HMMER3 profile HMMs (70) from the EggNOG database (76) for the proteins and Infernal covariance models (69) from the Rfam database (73) for the RNAs. Missing values due to method failure were again recorded, and in the worst case affected 1.5% of the simulations, again, not enough to alter our main conclusion. The results can be viewed in Figure 2A&B and Supplementary Tables 3&6. The ΔΔ G and Δ bitscore for proteins and RNAs were only compared qualitatively, since the methods and models to compute each are quite different.

The corresponding computer code for the above steps is available in `computeDeltaDelta.pl` and `plotDelta.R` scripts in the github repository. Raw data is available in the `data/delta-delta/` directory.

The RNA sequences of SgrS RNA (Rfam accession: RF00534, 227 nucleotides long) and corresponding protein SgrT (Pfam accession: PF15894, 102 nucleotides long) were mutated *in silico* with random substitution or indel mutations 100 times, with mutation rates varying between 0 and 500 per kilobase. We used `PSSpred` to infer the probability of each residue in the SgrT mutants forming an alpha helix, beta sheet or coil (79, 80). For the RNA sequences, we used “`RNAfold -p`”, an implementation of McCaskill’s RNA partition folding function (81) found in the Vienna RNA Package (75). The results can be viewed in Figure 2C&D.

The corresponding computer code for the above steps is available in `structureMutagenerator.pl` and `plotStructureMutagenerator.R` scripts in the github repository. Raw data is available in the `data/sgr-structural/` directory.

### Fluorescent protein and RNA construction and measurements

The fluorescent protein vector was constructed by inserting the mCherry gene into the NcoI and PmeI sites of pBAD-TOPO/LacZ/V5-His (Invitrogen) deriving pMCH01 (P_BAD_-mCherry, pBR322+ROP backbone, Amp^R^). Plasmid pBRC01 (T7-Broccoli-Broccoli, pBR322+ROP backbone, Kan^R^) was purchased as pET28c-F30-2xdBroccoli (Addgene) (Figure S5). Mutagenesis libraries were constructed using GeneMorph II Random Mutagenesis Kit (Agilent Technologies). The mCherry gene and Broccoli aptamer were amplified from their respective plasmids using Mutazyme II DNA polymerase to generate mega primers for MEGAWHOP whole plasmid PCR (82). Parental plasmids were digested with restriction enzyme DpnI, and the resulting mutation library was introduced into competent *E. coli* BL21(DE3) (Broccoli) or *E. coli* BL21(DE3) pLys (mCherry). We constructed two mCherry libraries with mutation rates of approximately one and four mutations per kilobase, and three Broccoli libraries with mutation rates of four, five and six mutation per kilobase. Approximately 10 clones from each library were sequenced to determine the mutation frequencies and whether the mutations were indels, transitions or transversions. Individual clones (*n* = 96) from each library were frozen for later analyses.

Cultures were grown at 37°C in Luria Bertani broth supplemented with appropriate antibiotics in a dry shaking incubator at 150 rpm. Each library was grown overnight in a 96-well plate before transfer to a second plate containing fresh medium supplemented with 1 mM isopropyl β-D-1-thiogalactopyranoside (IPTG) and 200 μM DHFB-T1 (Lucerna) to induce expression of Broccoli or 0.2% arabinose to induce expression of mCherry. We also prepared a plate containing eight wells of induced parental constructs (positive), uninduced parental constructs (negative), and LB supplemented with inducers (blank) for controls. The next morning, each library plate was used to culture three independent replica plates (three total cultures per mutant) and the control plate was used to culture one replica plate (eight total cultures per control condition). All plates were grown for 6 h before a Fluostar Omega plate reader (BMG Labtech) was used to measure the optical density (600 nm) and fluorescence. Fluorescence for the mCherry mutant library was measured with a 584 nm excitation filter and a 620 nm emission filter, with a 1500 gain. Fluorescence for the Broccoli mutant library was measured with a 485 nm excitation filter and a 520 nm emission filter, with a 1000 gain. Relative fluorescent units (RFU) was divided by optical density to derive a “Growth modified RFU”, and then by no-mutant controls to get the “Relative Fluorescence”. The no-mutant controls for the libraries were the parental plasmids and the no-mutant controls for the individual clones were unmutated clones within the library. A summary of the results can be viewed in Figure 3A-C.

The corresponding computer code for the above steps is available in `plotFluoro.R` script in the github repository. Raw data is available in the `data/fluoro/` directory.

## Classification

BIOLOGICAL SCIENCES: Biophysics and Computational Biology; Evolution; Genetics; Systems Biology.

## Data and software availability

The software, documentation, sequences, and results for this project are available on our github repository: https://github.com/Gardner-BinfLab/robustness-RNP.

## Acknowledgements

We thank the reviewer who challenged the senior author’s unsupported statement “RNAs, unlike proteins, are relatively robust to genetic variation” in a 2014 draft manuscript, thus inspiring this study. We thank Gabrielle David, PhD, for editing a draft of this manuscript. We are grateful to Dr Elena Rivas and Professor Dan Tawfik who gave critical feedback on the draft manuscript, and to the attendees of the 2018 Benasque “Computational Approaches to RNA Structure and Function” conference for many valuable discussions. This work was supported by Biomolecular Interaction Centre postdoctoral grants (DS Coray) and by a Rutherford Discovery Fellowship administered by the Royal Society of New Zealand awarded to P. Gardner.

## Notes

#### Summary of Updates

Substantially updated experiment 2: simulated mutagenesis, and the predicted impact on protein and RNA structure.

https://github.com/Gardner-BinfLab/robustness-RNP

## References

1. L. S. Waters, G. Storz, Regulatory RNAs in bacteria. Cell 136, 615–628 (2009).

2. T. R. Cech, J. A. Steitz, The noncoding RNA revolution-trashing old rules to forge new ones. Cell 157, 77–94 (2014).

3. W. Gilbert, Origin of life: The RNA world. Nature 319 (1986).

4. S. Lindgreen, et al., Robust identification of noncoding RNA from transcriptomes requires phylogenetically-informed sampling. PLoS Comput. Biol. 10, e1003907 (2014).

5. G.-S. Wang, T. A. Cooper, Splicing in disease: disruption of the splicing code and the decoding machinery. Nat. Rev. Genet. 8, 749–761 (2007).

6. J.-Q. Chen, et al., Variation in the ratio of nucleotide substitution and indel rates across genomes in mammals and bacteria. Mol. Biol. Evol. 26, 1523–1531 (2009).

7. J. A. G. M. de Visser, et al., Perspective: Evolution and detection of genetic robustness. Evolution 57, 1959–1972 (2003).

8. E. van Nimwegen, J. P. Crutchfield, M. Huynen, Neutral evolution of mutational robustness. Proceedings of the National Academy of Sciences 96, 9716–9720 (1999).

9. N. K. Mimouni, R. B. Lyngsø, S. Griffiths-Jones, J. Hein, An analysis of structural influences on selection in RNA genes. Mol. Biol. Evol. 26, 209–216 (2009).

10. C. Li, W. Qian, C. J. Maclean, J. Zhang, The fitness landscape of a tRNA gene. Science 352, 837–840 (2016).

11. G. Abrusán, J. A. Marsh, Alpha Helices Are More Robust to Mutations than Beta Strands. PLoS Comput. Biol. 12, e1005242 (2016).

12. H. H. Guo, J. Choe, L. A. Loeb, Protein tolerance to random amino acid change. Proc. Natl. Acad. Sci. U. S. A. 101, 9205–9210 (2004).

13. D. A. Drummond, J. D. Bloom, C. Adami, C. O. Wilke, F. H. Arnold, Why highly expressed proteins evolve slowly. Proc. Natl. Acad. Sci. U. S. A. 102, 14338–14343 (2005).

14. J. Stombaugh, C. L. Zirbel, E. Westhof, N. B. Leontis, Frequency and isostericity of RNA base pairs. Nucleic Acids Res. 37, 2294–2312 (2009).

15. A. L. Goldberg, R. E. Wittes, Genetic Code: Aspects of Organization. Science 153, 420–424 (1966).

16. S. Alkatib, et al., The contributions of wobbling and superwobbling to the reading of the genetic code. PLoS Genet. 8, e1003076 (2012).

17. M. O. Dayhoff, R. M. Schwartz, B. C. Orcutt, 22 a model of evolutionary change in proteins. Atlas of protein sequence and structure, 345–352 (1978).

18. D. Haig, L. D. Hurst, A quantitative measure of error minimization in the genetic code. J. Mol. Evol. 33, 412–417 (1991).

19. A. S. Novozhilov, Y. I. Wolf, E. V. Koonin, Evolution of the genetic code: partial optimization of a random code for robustness to translation error in a rugged fitness landscape. Biol. Direct 2, 24 (2007).

20. A. Wagner, Robustness and Evolvability in Living Systems (2013).

21. R. Geyer, A. Madany Mamlouk, On the efficiency of the genetic code after frameshift mutations. PeerJ 6, e4825 (2018).

22. L. Bartonek, D. Braun, B. Zagrovic, Invariants of Frameshifted Variants. bioRxiv, 684076 (2019).

23. L. E. Maquat, Defects in RNA splicing and the consequence of shortened translational reading frames. Am. J. Hum. Genet. 59, 279–286 (1996).

24. D. G. MacArthur, et al., Guidelines for investigating causality of sequence variants in human disease. Nature 508, 469–476 (2014).

25. T. Ohta, Slightly deleterious mutant substitutions in evolution. Nature 246, 96–98 (1973).

26. D. K. Chiu, T. Kolodziejczak, Inferring consensus structure from nucleic acid sequences. Comput. Appl. Biosci. 7, 347–352 (1991).

27. S. Henikoff, J. G. Henikoff, Amino acid substitution matrices from protein blocks. Proc. Natl. Acad. Sci. U. S. A. 89, 10915–10919 (1992).

28. A. Kun, M. Santos, E. Szathmáry, Real ribozymes suggest a relaxed error threshold. Nat. Genet. 37, 1008–1011 (2005).

29. A. Babajide, I. L. Hofacker, M. J. Sippl, P. F. Stadler, Neutral networks in protein space: a computational study based on knowledge-based potentials of mean force. Fold. Des. 2, 261–269 (1997).

30. E. Ferrada, A. Wagner, A Comparison of Genotype-Phenotype Maps for RNA and Proteins. Biophys. J. 102, 1916–1925 (2012).

31. S. F. Greenbury, S. Schaper, S. E. Ahnert, A. A. Louis, Genetic Correlations Greatly Increase Mutational Robustness and Can Both Reduce and Enhance Evolvability. PLoS Comput. Biol. 12, e1004773 (2016).

32. S. E. Ahnert, Structural properties of genotype-phenotype maps. J. R. Soc. Interface 14 (2017).

33. M. You, S. R. Jaffrey, Structure and Mechanism of RNA Mimics of Green Fluorescent Protein. Annu. Rev. Biophys. 44, 187–206 (2015).

34. G. S. Filonov, J. D. Moon, N. Svensen, S. R. Jaffrey, Broccoli: rapid selection of an RNA mimic of green fluorescent protein by fluorescence-based selection and directed evolution. J. Am. Chem. Soc. 136, 16299–16308 (2014).

35. O. Shimomura, F. H. Johnson, Y. Saiga, Extraction, purification and properties of aequorin, a bioluminescent protein from the luminous hydromedusan, Aequorea. J. Cell. Comp. Physiol. 59, 223–239 (1962).

36. F. G. Prendergast, K. G. Mann, Chemical and physical properties of aequorin and the green fluorescent protein isolated from Aequorea forskalea. Biochemistry 17, 3448–3453 (1978).

37. R. Y. Tsien, The green fluorescent protein. Annu. Rev. Biochem. 67, 509–544 (1998).

38. J. Felsenstein, DNAML in PHYLIP 2.6. University of Washington, Seattle (1984).

39. R. D. Finn, et al., The Pfam protein families database: towards a more sustainable future. Nucleic Acids Res. 44, D279–85 (2016).

40. I. Kalvari, et al., Rfam 13.0: shifting to a genome-centric resource for non-coding RNA families. Nucleic Acids Res. 46, D335–D342 (2018).

41. M. Long, E. Betrán, K. Thornton, W. Wang, The origin of new genes: glimpses from the young and old. Nat. Rev. Genet. 4, 865–875 (2003).

42. B. R. Jose, P. P. Gardner, L. Barquist, Transcriptional noise and exaptation as sources for bacterial sRNAs. Biochem. Soc. Trans. 47, 527–539 (2019).

43. H. B. Fraser, A. E. Hirsh, L. M. Steinmetz, C. Scharfe, M. W. Feldman, Evolutionary rate in the protein interaction network. Science 296, 750–752 (2002).

44. H. M. Berman, et al., The Protein Data Bank. Nucleic Acids Res. 28, 235–242 (2000).

45. N. Tokuriki, F. Stricher, J. Schymkowitz, L. Serrano, D. S. Tawfik, The stability effects of protein mutations appear to be universally distributed. J. Mol. Biol. 369, 1318–1332 (2007).

46. J. Ritz, J. S. Martin, A. Laederach, Evaluating our ability to predict the structural disruption of RNA by SNPs. BMC Genomics 13 Suppl 4, S6 (2012).

47. N. E. Wheeler, L. Barquist, R. A. Kingsley, P. P. Gardner, A profile-based method for identifying functional divergence of orthologous genes in bacterial genomes. Bioinformatics 32, 3566–3574 (2016).

48. N. E. Wheeler, P. P. Gardner, L. Barquist, Machine learning identifies signatures of host adaptation in the bacterial pathogen Salmonella enterica. PLoS Genet. 14, e1007333 (2018).

49. M. Halvorsen, J. S. Martin, S. Broadaway, A. Laederach, Disease-Associated Mutations That Alter the RNA Structural Ensemble. PLoS Genet. 6, e1001074+ (2010).

50. J. S. Mattick, I. V. Makunin, Non-coding RNA. Hum. Mol. Genet. 15 Spec No 1, R17–29 (2006).

51. R. Hershberg, S. Altuvia, H. Margalit, A survey of small RNA-encoding genes in Escherichia coli. Nucleic Acids Res. 31, 1813–1820 (2003).

52. B. Rost, Twilight zone of protein sequence alignments. Protein Eng. 12, 85–94 (1999).

53. E. K. Freyhult, J. P. Bollback, P. P. Gardner, Exploring genomic dark matter: a critical assessment of the performance of homology search methods on noncoding RNA. Genome Res. 17, 117–125 (2007).

54. E. P. Nawrocki, S. R. Eddy, Query-dependent banding (QDB) for faster RNA similarity searches. PLoS Comput. Biol. 3, e56 (2007).

55. S. R. Eddy, R. Durbin, RNA sequence analysis using covariance models. Nucleic Acids Res. 22, 2079–2088 (1994).

56. P. A. G. Tizei, E. Csibra, L. Torres, V. B. Pinheiro, Selection platforms for directed evolution in synthetic biology. Biochem. Soc. Trans. 44, 1165–1175 (2016).

57. R. E. Lenski, Experimental evolution and the dynamics of adaptation and genome evolution in microbial populations. ISME J. 11, 2181–2194 (2017).

58. W. Fontana, P. Schuster, Continuity in evolution: on the nature of transitions. Science 280, 1451–1455 (1998).

59. V. Moulton, et al., RNA folding argues against a hot-start origin of life. J. Mol. Evol. 51, 416–421 (2000).

60. C. P. Lepper, M. A. K. Williams, P. J. B. Edwards, V. V. Filichev, G. B. Jameson, Effects of Pressure and pH on the Physical Stability of an I-Motif DNA Structure. ChemPhysChem 20, 1567–1571 (2019).

61. R. C. McBride, C. B. Ogbunugafor, P. E. Turner, Robustness promotes evolvability of thermotolerance in an RNA virus. BMC Evol. Biol. 8, 231 (2008).

62. R. E. Lenski, J. E. Barrick, C. Ofria, Balancing robustness and evolvability. PLoS Biol. 4, e428 (2006).

63. J. D. Bloom, S. T. Labthavikul, C. R. Otey, F. H. Arnold, Protein stability promotes evolvability. Proc. Natl. Acad. Sci. U. S. A. 103, 5869–5874 (2006).

64. A. M. Poole, D. C. Jeffares, D. Penny, The path from the RNA world. J. Mol. Evol. 46, 1–17 (1998).

65. M. Madera, J. Gough, A comparison of profile hidden Markov model procedures for remote homology detection. Nucleic Acids Res. 30, 4321–4328 (2002).

66. Z. Li, Y. Zhang, Predicting the secondary structures and tertiary interactions of 211 group I introns in IE subgroup. Nucleic Acids Res. 33, 2118–2128 (2005).

67. T. M. Lowe, P. P. Chan, tRNAscan-SE On-line: integrating search and context for analysis of transfer RNA genes. Nucleic Acids Res. 44, W54–7 (2016).

68. I. Kalvari, et al., Rfam 13.0: shifting to a genome-centric resource for non-coding RNA families. Nucleic Acids Res. 46, D335–D342 (2018).

69. E. P. Nawrocki, S. R. Eddy, Infernal 1.1: 100-fold faster RNA homology searches. Bioinformatics 29, 2933–2935 (2013).

70. S. R. Eddy, Accelerated Profile HMM Searches. PLoS Comput. Biol. 7, e1002195 (2011).

71. M. Suyama, D. Torrents, P. Bork, PAL2NAL: robust conversion of protein sequence alignments into the corresponding codon alignments. Nucleic Acids Res. 34, W609–12 (2006).

72. H. B. Mann, D. R. Whitney, On a Test of Whether one of Two Random Variables is Stochastically Larger than the Other. Ann. Math. Stat. 18, 50–60 (1947).

73. S. R. Eddy, A. Bateman, R. D. Finn, A. I. Petrov, Rfam 13.0: shifting to a genome-centric resource for non-coding RNA families. Nucleic acids (2017).

74. J. Felsenstein, PHYLIP version 3.6. Software package, Department of Genome Sciences, University of Washington, Seattle, USA (2005).

75. R. Lorenz, et al., ViennaRNA Package 2.0. Algorithms Mol. Biol. 6, 26 (2011).

76. J. Huerta-Cepas, et al., eggNOG 4.5: a hierarchical orthology framework with improved functional annotations for eukaryotic, prokaryotic and viral sequences. Nucleic Acids Res. 44, D286–93 (2016).

77. J. Laimer, H. Hofer, M. Fritz, S. Wegenkittl, P. Lackner, MAESTRO--multi agent stability prediction upon point mutations. BMC Bioinformatics 16, 116 (2015).

78. I. L. Hofacker, et al., Fast folding and comparison of RNA secondary structures. Monatshefte für Chemie / Chemical Monthly 125, 167–188 (1994).

79. R. Yan, D. Xu, J. Yang, S. Walker, Y. Zhang, A comparative assessment and analysis of 20 representative sequence alignment methods for protein structure prediction. Sci. Rep. 3, 2619 (2013).

80. J. Yang, et al., The I-TASSER Suite: protein structure and function prediction. Nat. Methods 12, 7–8 (2015).

81. J. S. McCaskill, The equilibrium partition function and base pair binding probabilities for RNA secondary structure. Biopolymers 29, 1105–1119 (1990).

82. K. Miyazaki, MEGAWHOP cloning: a method of creating random mutagenesis libraries via megaprimer PCR of whole plasmids. Methods Enzymol. 498, 399–406 (2011).

