## Supplementary figures for "The genetic robustness of RNA and protein from evolutionary, structural and functional perspectives"

### Supplementary Material

**Supplementary Tables:**

**Table S1: Bacterial RNPs and data for Figure 1A**
**Table S2: Fungal RNPs and data for Figure 1B**
**Table S3: RNPs for structural analysis for Figure 2A&B**
**Table S4: Fluorescence data for Figure 3**
**Table S5: Fluorescent RNA and protein sequences for Figure 3**
https://docs.google.com/spreadsheets/d/1exZaYpTQRfTpdNBVaIOJID3Uzw_WIJy0XBOTmPaOSi4/edit?usp=sharing

#### Comparison of neutral variation between extant sequences in RNPs

Variant sites identified between deeply or recently conserved RNPs (Table S1&S2) were identified and classified as neutral if it preserved structure and/or biochemistry (Figure S1A&B). In RNA secondary structure was preserved more than biochemistry (Figure S1C). In protein on average a third to two-thirds (~0.3 bacteria, ~0.6 fungi) of the variations were synonymous, with many of the non-synonymous variations preserving biochemistry of the coded amino acid according to two measures (Figure S2C). Bacterial protein and RNA had higher proportions of non-neutral variation than fungi.

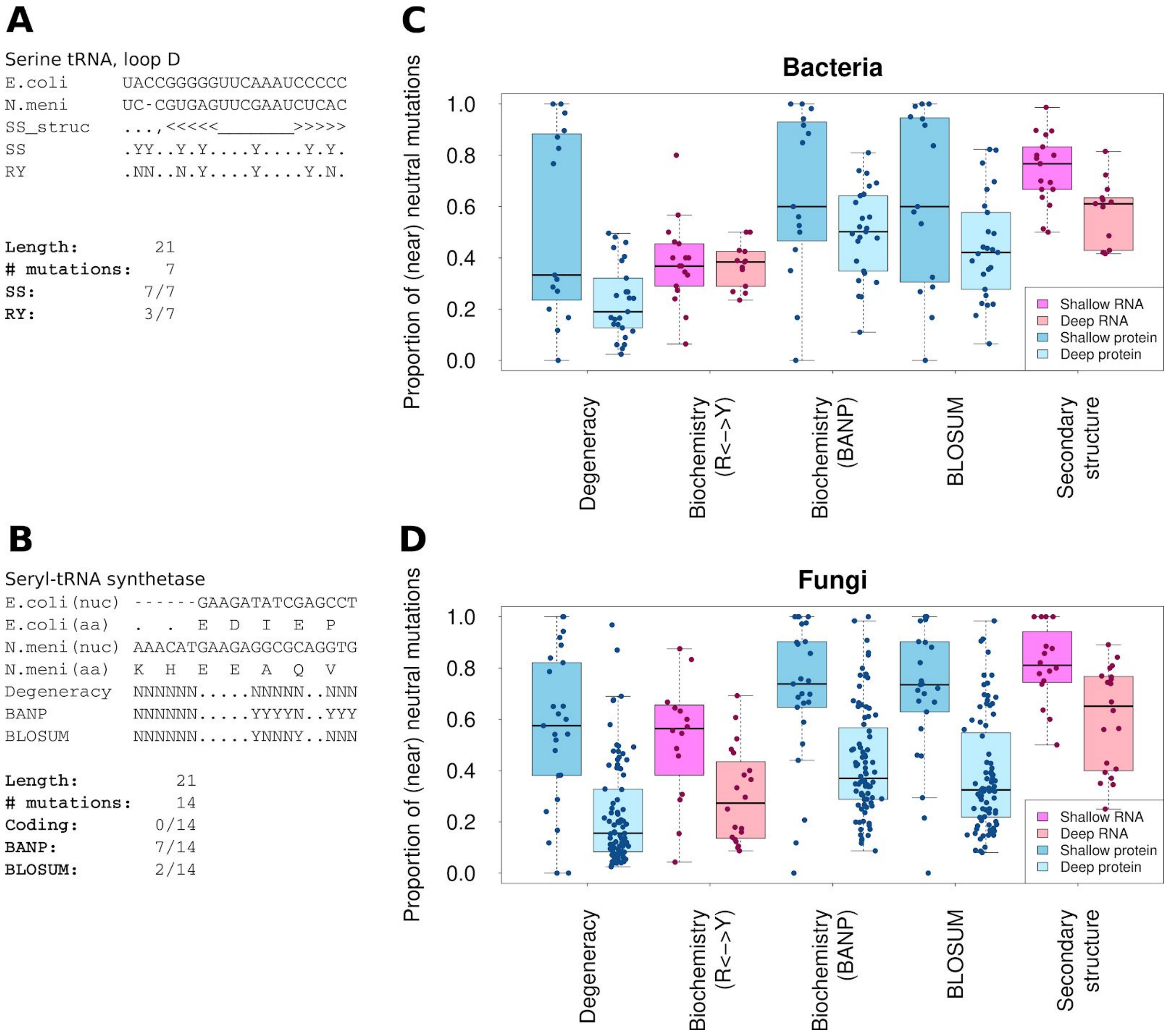


**Figure S1: The proportion of nucleotide variants in RNA or protein that can be classed as neutral.** A collection of functionally linked RNA and protein families that are shared between *E. coli* and *N. meningitidis* (*N. meni*) (Deep, lighter shades) or between *E. coli* and *S. enterica* (Shallow, darker shades). Each nucleotide variant is classified as either neutral or non-neutral according to a number of different models. (**A&B**) Exemplar genome variants and different classification schemes. (**A**) To score differences in the RNA serine tRNA, for example, secondary structure of each was determined (**SS_struct**) and changes between species (in red) was scored as either near-neutral or not, for changes in secondary structure (**SS** or **Secondary structure**) or biochemistry (**RY** show transitions, R: A<->G, Y: C<->U). (**B**) To score differences in the protein seryl-tRNA synthetase. For example, both nucleotide (nuc) and amino acid (aa) sequences were compared across the two species. The nucleotide differences between species was scored as neutral if the resulting amino acids were the same, labelled **Degeneracy**. Biochemically neutral variation, labelled **BANP,** classed the following groups of replacements as neutral (**B**asic (H,R,K), **A**cidic (D,E), **N**on-polar (F,L,W,P,I,M,V,A) or **P**olar (G,S,Y,C,T,N,Q)) or if amino acid replacements were assigned a non-negative score in the **BLOSUM** score matrix (27). (**C&D**) The proportion of near-neutral mutations for each RNA or protein was compared for different models of neutrality across deep and shallow phylogenetic distances for RNAs and proteins from bacteria (**C**) or fungi (**D**). The x-axis labels are described above.

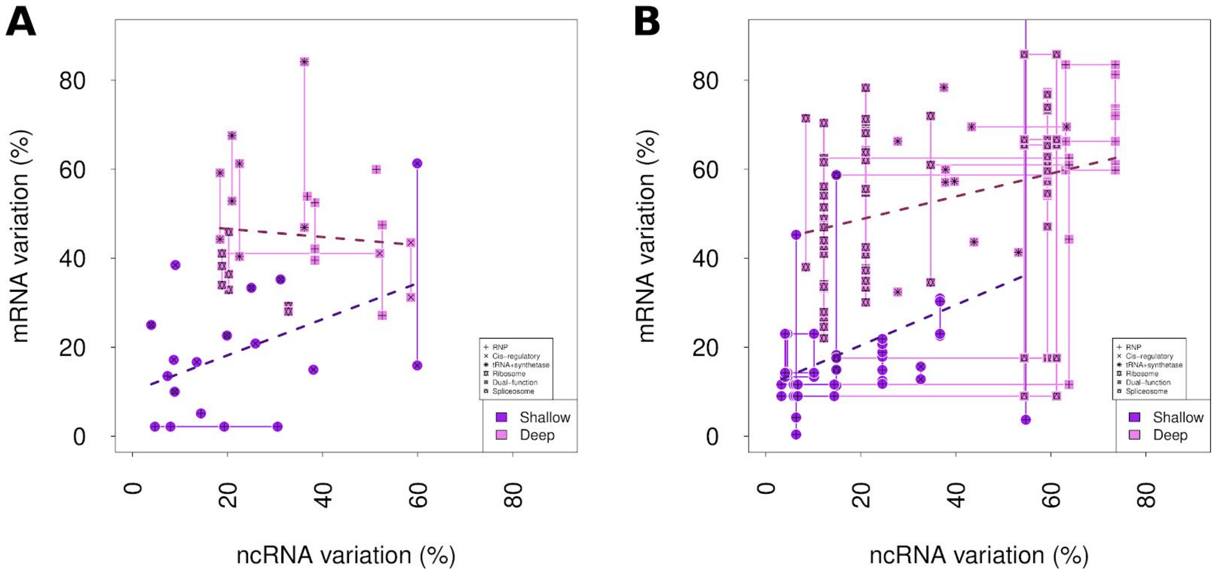


**Figure S2: Level of nucleotide variation interacting RNA and protein pairs.** Interacting RNA and protein pairs were compared across shallow and deep timescales in bacteria (**A**) and fungi (**B**). The percent nucleotide variation each RNA-protein pair was determined, for either shallow or deep divergence times. There was little correlation (Spearman's) between the variation in one molecule and its interacting partner in either the bacteria deep (*Rho* = -0.04, *p* = 0.85) or shallow (*Rho* = 0.29, *p* = 0.24) or fungi deep (*Rho* = 0.37, *p* = 0.0001 ) or shallow (*Rho* = 0.26 , *p* = 0.07 ) groups.

#### Maintenance of structure after in *silico* mutagenesis of RNPs

**Table S6:** Columns 1-6 contain the proportions of simulated point mutations in the protein and RNA components of RNPs from the PDB that can be classified as destabilising, divergent or both using estimated $\Delta\Delta G$ and $\Delta$bitscore values (see Figure 2A&B). Columns 7 and 8 contain the Spearman correlation coefficients and corresponding P-values for the relationships between the protein and RNA $\Delta\Delta G$ and $\Delta$bitscore values (see also Figure S3). In each case there is a significant relationship between the two, except for proteins with a single variant site.

| | **Destabilising ($\Delta\Delta G$>0)** | | **Divergent ($\Delta$bitscore<0)** | | **Destabilising & Divergent ($\Delta\Delta G$>0 & $\Delta$bitscore<0)** | | **Spearman correlation coefficient (P-value)** | |
|---|---|---|---|---|---|---|---|---|
| | **Protein** | **RNA** | **Protein** | **RNA** | **Protein** | **RNA** | **Protein** | **RNA** |
| **One mutn** | 30% | 41% | 50% | 61% | 60% | 66% | -0.01 (0.67) | -0.35 ($<2.2e^{-16}$) |
| **Four mutns** | 58% | 73% | 89% | 96% | 98% | 93% | -0.20 ($<2.2e^{-16}$) | -0.25 ($<2.2e^{-16}$) |

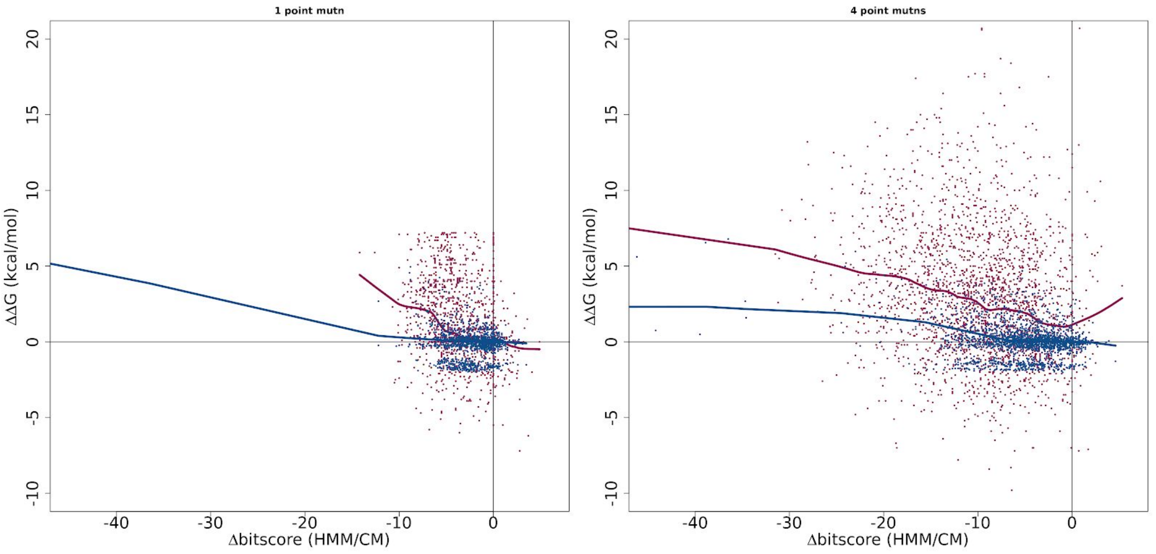


**Figure S3: Relationship between $\Delta\Delta G$ and $\Delta$bitscore values for simulated protein and RNA mutations.** Random mutations were introduced into mRNAs (blue) or ncRNAs (pink) for RNPs with solved structures. $\Delta\Delta G$ and $\Delta$bitscore values were computed as described in the methods, and have been used as an estimate of the impact of random simulated mutations on protein and RNA structures.

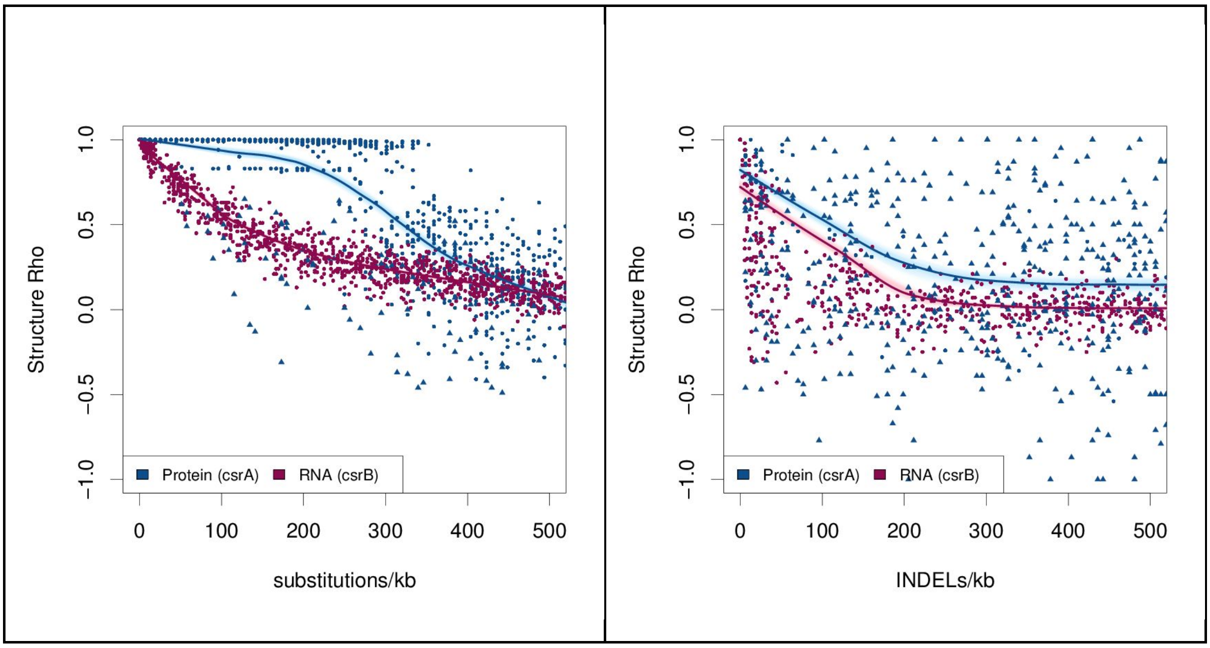


**Figure S4: Structural robustness of the CsrA protein and CsrB sRNA.** Random mutants of the CsrA messenger RNA (blue) and CsrB small RNA (pink) were generated *in silico*. Their secondary structure probabilities were predicted using "RNAfold-p" and "PSSpred". The per-residue probabilities of either base-paired/not-base-paired or alpha/beta/coil were compared between native and mutated sequences using Spearman's correlation. This gave a "structure rho", where 1 implies the predicted mutant structure is identical to the predicted parental structure, 0 means there is no correlation, and −1 shows a perfect inverse correlation. (A) Substitution mutations and (B) insertion or deletion mutations (indels) were introduced into the RNA (pink) and protein (blue) at rates ranging from 1 to 500 mutations per kilobase (kb). Points corresponding to truncated protein or small RNA with a length less than 75% that of the original are indicated with a solid triangle, otherwise a solid circle is used. Local polynomial regression (loess) curves were fitted to the RNA and protein points. To indicate the confidence for each loess curve, these were bootstrapped 500 times and plotted in light pink or blue to resampled points.

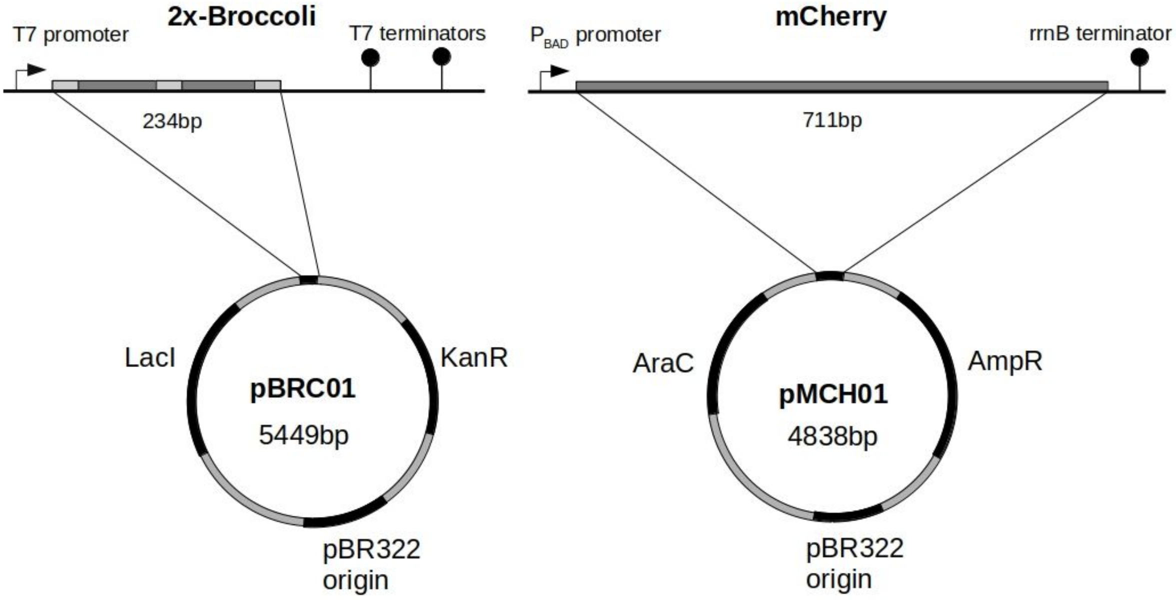


**Figure S5: Parental plasmids for whole-plasmid PCR mutagenesis of mCherry and Broccoli**
Plasmid pBRC01 contains a 2x-Broccoli construct behind a T7 promoter made up of two broccoli aptamers (dark grey) between an F30 (light grey) scaffold that aids in folding, followed by two T7 promoters. The vector also includes a LacI control protein for expression of the chromosomal T7 polymerase and a Kanamycin resistance gene. Plasmid pMCH01 contains mCherry construct behind a $P_{BAD}$ promoter and followed by an rrnB terminator. The vector also includes an AraC protein for control of the $P_{BAD}$ promoter and an Ampicillin resistance gene. Both are on mid-copy plasmid pBR322+ROP.

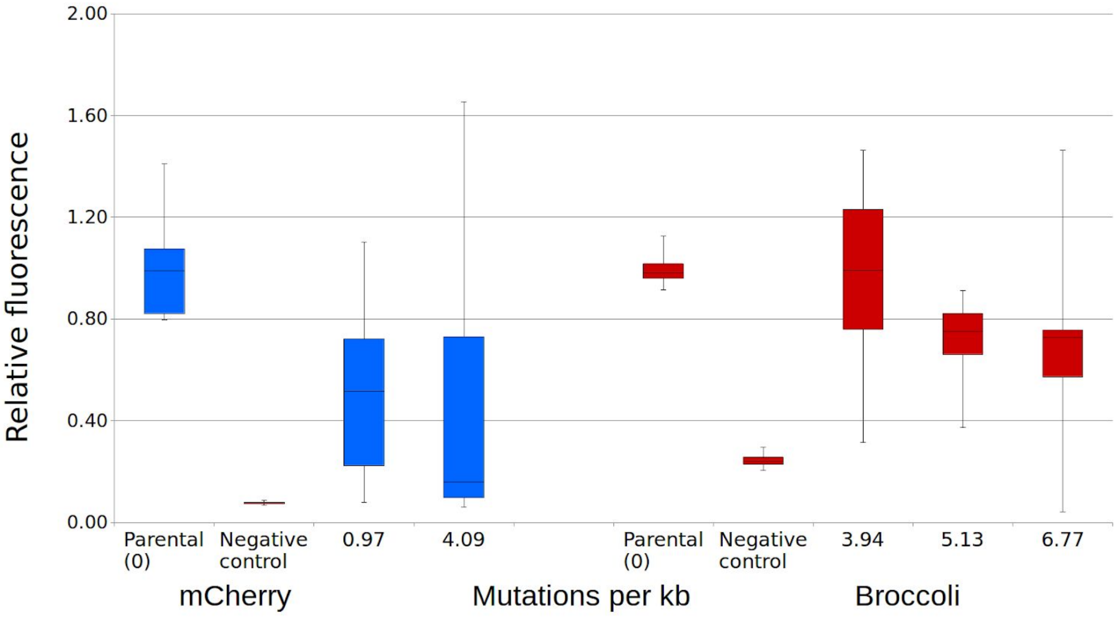


**Figure S6: Fluorescence of mutant libraries of RNA aptamer Broccoli and protein mCherry.** Libraries of randomly mutated fluorescent RNA aptamer Broccoli and fluorescent protein mCherry were tested for function relative to the unmutated control. Two libraries of mCherry and three libraries of Broccoli were constructed, using a range of mutation rates per kilobase (kb). The fluorescence intensities of the mutants were normalized to the optical density and the fluorescence intensities of unmutated controls.
